# MINI-EX version 2: cell-type-specific gene regulatory network inference using an integrative single-cell transcriptomics approach

**DOI:** 10.1101/2023.12.24.573246

**Authors:** Jasper Staut, Nicolás Manosalva Pérez, Thomas Depuydt, Klaas Vandepoele, Svitlana Lukicheva

## Abstract

Understanding and predicting cell-type-specific gene regulatory networks (GRNs) is essential for unraveling the complex interactions between transcription factors (TFs) that modulate the expression of target genes and control diverse biological processes in multicellular organisms. MINI-EX (Motif-Informed Network Inference based on single-cell EXpression data) is an integrative tool tailored for identifying cell-type-specific GRNs in plants. Leveraging single-cell transcriptomics data, MINI-EX constructs expression-based networks and integrates TF motif information to produce GRNs with increased accuracy. Furthermore, it assigns regulatory modules to distinct cell types and prioritizes candidate regulators by employing a strategy that encompasses network centrality measures, functional annotations, and expression specificity. Taken together, MINI-EX offers a powerful approach to identify cell-type-specific transcriptional cascades and enhance our understanding of TF functions in plant biology. Here, we discuss recent advancements in the tool’s latest version and explain how single-cell GRNs can be identified for non-model species lacking TF motif information. Additionally, we provide a comprehensive guide to use MINI-EX, covering the entire pipeline from preparing input files starting from a single-cell experiment, over configuring parameters, to interpreting output data.

## 1. Introduction

Recently, single-cell technologies have been increasingly used to gain insight into complex biological mechanisms (***1–4***). Zooming in on the level of the single cell allows tracking dynamic cellular processes, study rare cell populations, and differentiate between tissue-specific responses to stress or hormone treatment experiments (***1, 5***). While initial single-cell studies in plants focused on model plant *Arabidopsis thaliana* (***4, 6–10***), the plant single-cell field has since been extended to important crop species such as maize, rice and tomato (***11–14***). With the new opportunities the single-cell technology brings, also comes the computational challenge to represent, integrate and leverage the information these datasets encompass, with the ultimate goal to extract novel, biological knowledge from them. With this objective in mind, gene regulatory network (GRN) inference can offer a suitable approach to integrate data and represent the regulatory relationships between genes, facilitating the identification of key regulators (***15–17***). The GRNs consist of nodes, representing genes, and directed edges between them, denoting a regulatory relationship between a transcription factor (TF) to a target gene (TG) (***18***). TFs are proteins that bind to specific DNA sequences, called motifs, to regulate the expression of a TG. Together, one TF and its TGs are referred to as a regulon.

In a recent publication, MINI-EX, a tool for cell-type-specific GRN inference and prioritization of candidate regulators in plants was presented (***19***). It is designed to integrate single-cell RNA sequencing data (scRNA-Seq) with TF binding motif information, inferring high-quality GRNs for each cell type and prioritize most relevant regulators using network statistics and gene ontology (GO; a controlled vocabulary describing current knowledge about the function and location of genes products) information. Both in its initial release and subsequent comparisons, MINI-EX has been shown to outperform the previously developed GRN inference tool SCENIC and is gaining interest within the plant single-cell community (***19–21***). In the recent release of MINI-EX version 2.0, its algorithm, visualizations and documentation have been further improved. In addition, new features were added such as the possibility to omit the integration of TF binding motif information, which provides a quick and straightforward way to run MINI-EX for non-supported species without the need for additional pre-processing of motif data for those species. In this chapter, we offer a practical guide on how to use MINI-EX for cell-type-specific GRN inference. We will cover the requirements for running the algorithm, briefly explain its basic principles, show how to prepare the configuration and input files, and demonstrate how to run the pipeline. Finally, we discuss all output files and provide an example of how the results from MINI-EX can be interpreted to advance understanding of biological processes.

## 2. Materials

### 2.1. Hardware and software

MINI-EX, implemented as a Nextflow (***22***) pipeline and distributed with a Singularity (***23***) container, can be run on any POSIX compatible system (such as Linux, OS X, …) and requires only Nextflow and Singularity installations for its execution. Since MINI-EX is designed to handle large datasets, it is recommended for use on high performance computing (HPC) clusters. Version 2.2 of MINI-EX (used in this chapter) requires approximately 17 hours to run on the Wendrich *et al.* (2020) dataset discussed in this chapter (and only 30 minutes when using a precomputed GRNBoost2 output file) and uses a maximum of 20 Gb of RAM (detailed memory requirements per process can be found in the config file available on GitHub). The example dataset provided on the GitHub repository can be run on a personal computer and takes around 18 minutes for the full run and around three minutes with a precomputed GRNBoost2 output file.

### 2.2. Expertise

MINI-EX execution begins with information extracted from a processed single-cell RNA sequencing (scRNA-seq) dataset. Therefore, we assume that the user possesses the capability to conduct a conventional single-cell processing analysis. Further in the chapter, we provide detailed instructions on how to extract MINI-EX input files from a Seurat object; an example of processing scRNA-Seq data using the Seurat library can be found in (***24***). However, users are free to use alternative tools, provided they can extract the required MINI-EX input files in the correct format.

### 2.3. Input files

To execute MINI-EX, the four following files, derived from processing an experimental scRNA-Seq dataset, are required:

1. an expression matrix, which provides the expression levels of each gene (row) in each cell (column);
2. a list of marker genes per cluster, in Seurat’s findAllMarkers output format;
3. a file containing the cluster identity of each cell;
4. a file containing the cluster annotations.

MINI-EX currently provides support for four plant species: *Arabidopsis thaliana*, *Oryza sativa* (rice), *Zea mays* (mays) and *Solanum lycopersicum* (tomato). For these species, the user is able to fully utilize all MINI-EX features. For users interested in running MINI-EX on plant species not currently supported, it is possible to disable the motif analysis step (see Note 1). In this case, users only have to provide a list of TFs of the species of interest. Additional files may be generated (see Section 3.5 on how to do this) to leverage the full capabilities of MINI-EX, but are not required.

### 2.4. Example dataset

To illustrate the execution of MINI-EX and the evaluation of its output files, we will use the Wendrich Arabidopsis scRNA-Seq root dataset published in (***4***). The corresponding Seurat file is available for download from the MINI-EX GitHub page at the following link: https://github.com/VIB-PSB/MINI-EX/tree/main.

In summary, the input FASTQ files were initially processed using CellRanger v6.1.1. The resulting files were then employed to create the Seurat object using the Seurat R package v4.0.1. We set upper (2%) and lower (25%) thresholds on library size, merged the samples, and performed normalization using Seurat’s SCTransform function (***25***), with the number of variable features set to 7000. For UMAP creation, we used 70 principal components with a resolution of 1.2. After processing, this dataset contains 11,456 cells, 21,675 expressed genes, and 38 clusters, encompassing 15 tissues.

## 3. Methods

### 3.1. Basic principles

MINI-EX utilizes scRNA-seq data as input to infer cell-type-specific GRNs and prioritize relevant regulators. The MINI-EX algorithm consists of four general steps (Figure 1). In step 1, a coexpression-based GRN is generated using GRNBoost2 (***26***) yielding an initial expression-based network. In step 2, this network is filtered by retaining only putative TF-TG interactions if the TF binding motif is found to be enriched (through motif mapping and statistical analysis) in the regulatory region (i.e., 5000 bp upstream and 1000 bp downstream of the gene’s translation start/stop site and its introns) of a TG, and the regulon is enriched for the TF motif. This motif-filtered network is then used in step 3 where the expression of TFs and TGs in each cluster is used as a last filtering step, resulting in multiple cluster-specific GRNs. Finally, step 4 integrates network statistics and optional gene ontology (GO) information (functional annotations of genes) to rank the regulons in each cell-type on their importance.

**Figure 1.**
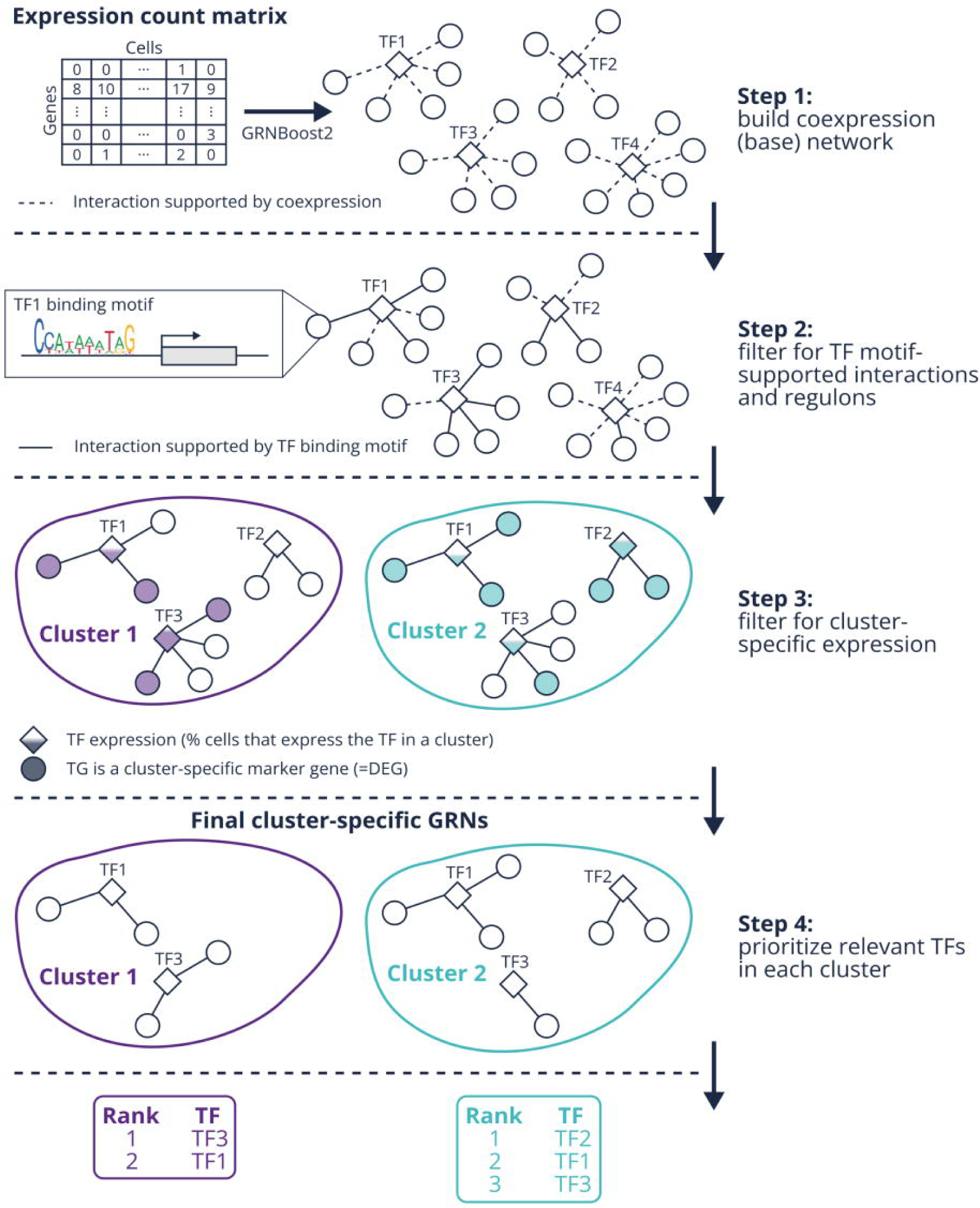
Generalized MINI-EX workflow. The main steps of the MINI-EX workflow are illustrated, showing a collection of regulons, each consisting of a transcription factor (TF; shown as diamonds) and its putative target genes (TGs; shown as circles), together forming a gene regulatory network (GRN). Steps 1 and 2 result in a single GRN for the full dataset that forms the precursor for step 3, which operates in a cluster-specific manner, yielding a unique GRN for each cluster of the single-cell input data. In step 4, TFs are prioritized based on network statistics and optional gene ontology terms (not shown).

### 3.2. MINI-EX 2.0 and later

In this section, we will briefly outline the changes that were made between the original MINI-EX release (version 1.0) and its later iterations. Given that this book chapter describes the use of MINI-EX 2.2, we want to clearly specify the differences between the original MINI-EX 1.0 and the version that is used here. The most relevant changes are listed below.

Version 2.0

1. MINI-EX can be easily run for any non-supported species by adding the possibility to omit the motif enrichment step (see step 2, Figure 1). Doing so lowers the precision of the resulting networks (see Note 1 for more details) but removes the need for pre-processed motif information that is only provided for supported species.
2. Support for tomato genome version SL4.0/ITAG4.0.
3. Addition of a new regulator heatmap visualization, called “regmap”. This visualization allows inspecting regulator importance over different cell-type clusters and/or developmental trajectories.
4. The procedure to calculate enrichment statistics is modified to conform with community guidelines (***27, 28***). With the aim of improving performance, the enrichment procedure is modified which results in differences in output between version 1.0 and 2.0 or later (see Note 2 for more details).
5. Other minor improvements are implemented such as bug fixes, improved documentation and input checks.

Version 2.1

6. Support for maize genome version AGPv5.
7. Addition of computationally inferred GO terms to the gene-GO association files (***29***) provided for supported species, except for *A. thaliana* (see Note 3 for more details).

Version 2.2

8. Added high-throughput evidence codes to the Arabidopsis gene-GO association file.
9. Improvements in the enrichment algorithm.
10. Added log file summarizing statistics on the input datasets and intermediate results in the workflow.
11. Improved layout of the generated figures.
12. Gene aliases file is made optional.

### 3.3. The MINI-EX configuration file

Before running MINI-EX, the user first needs to prepare the configuration (config) file, which contains all settings that the user needs to specify before starting the pipeline. This includes the path of all input files and the output folder, all parameters, as well as memory settings. In this section we describe the general setup of the config file. In Sections 3.4 and 3.5 we describe how to generate the input files and Section 3.6 covers the MINI-EX parameters. Changing the memory settings is covered in Section 3.7, which also explains how to run the pipeline.

The config file starts with two smaller sections. The first section specifies the job scheduler or executor and is covered in Section 3.7 about how to run the pipeline. The second section relates to the singularity container that MINI-EX uses to handle dependencies and should not be adapted by the user. The following section specifies input files and parameters. Figure 2 shows the MINI-EX algorithm workflow, including all input files and parameters that are to be set in the config file. Input files that are required by MINI-EX fall in two categories: input files to be provided by the user and files shipped with MINI-EX for supported species (see Figure 2). The user-provided files are specified first in the config file. These files are mandatory input and include four files derived from a single-cell dataset:

expressionMatrix = "$baseDir/example/INPUTS/*_matrix.tsv"
markersOut = "$baseDir/example/INPUTS/*_allMarkers.tsv"
cellsToClusters = "$baseDir/example/INPUTS/*_cells2clusters.tsv"
clustersToIdentities = "$baseDir/example/INPUTS/*_identities.tsv"

**Figure 2.**
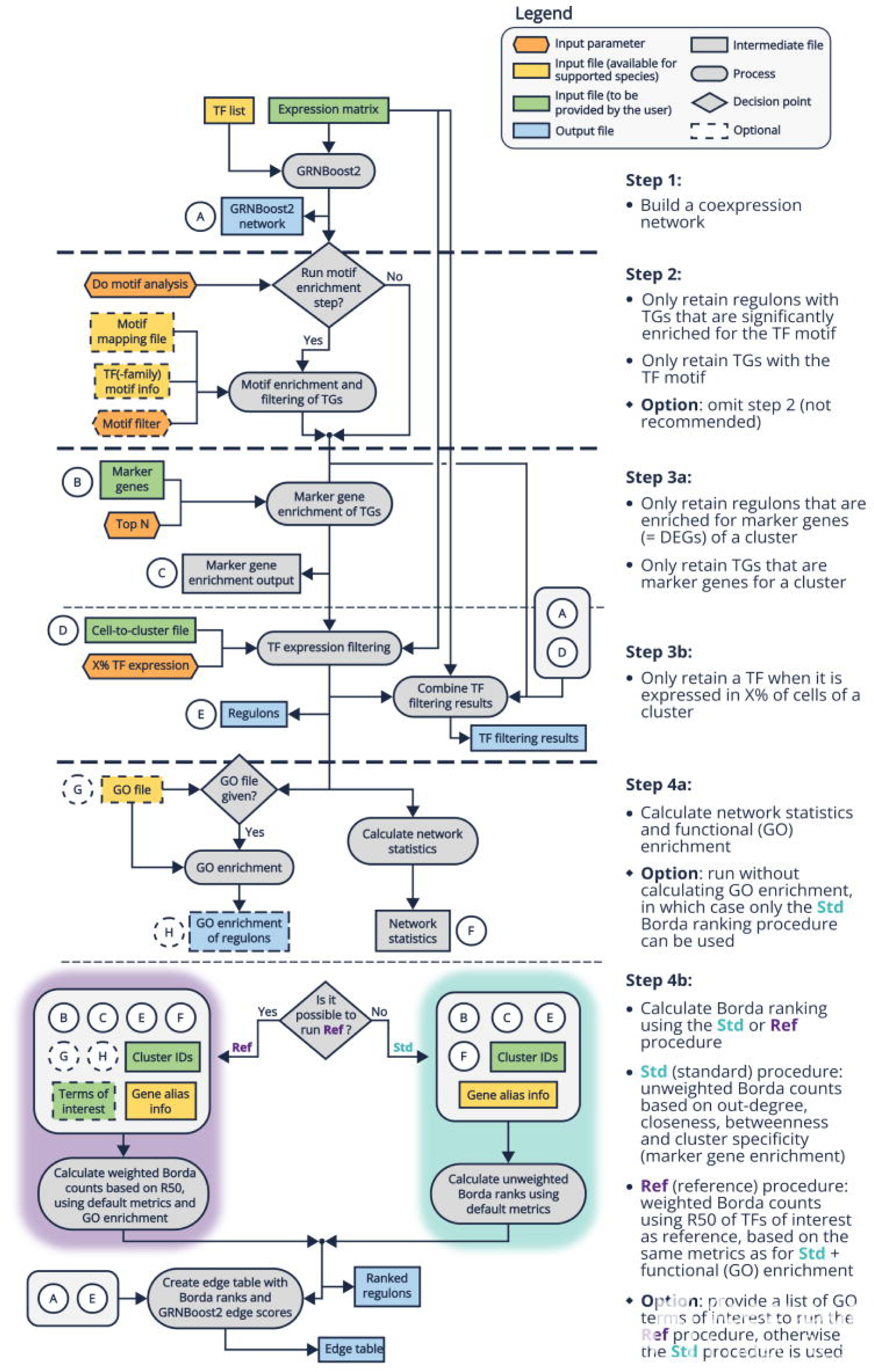
MINI-EX algorithm overview. Input files and parameters are used in a sequence of processes, producing several output files. Different options (shown as diamonds) are available for the user, corresponding with decision points in the workflow. Processes and files related to visualizations are not included in this schema.

The wildcard character ‘*’ in the file names above allows Nextflow to recognize input files with a different prefix (followed by an underscore) as coming from different datasets and run them in parallel (see Section 3.7). Files of the same dataset must always share the same prefix.

The second type of files includes generic data related to the species of interest. These files are available on the MINI-EX GitHub repository for the four supported species, but must be provided by the user for the non-supported species. In this case, only the list of transcription factors is mandatory; the remaining files should be specified if the user wants to fully utilize all of MINI-EX’s features:

tfList = "$baseDir/data/ath/ath_TF_list.tsv"
geneAliases = "$baseDir/data/ath/ath_gene_aliases.tsv"
infoTf = "$baseDir/data/ath/ath_TF2fam2mot.tsv"
featureFileMotifs = "$baseDir/data/ath/ath_2021.1_motifMapping.out.gz"
goFile = "$baseDir/data/ath/ath_full_BP_expcur_ext_names.tsv"

After the input files, parameters are defined:

doMotifAnalysis = true
termsOfInterest = "$baseDir/example/INPUTS/GOsIwant.txt"
motifFilter = "TF-F_motifs"
topMarkers = "700"
expressionFilter = "10"
topRegulons = "150"

The last section of the config file sets the memory for the different processes and will be discussed in Section 3.7.

### 3.4. Preparing input files from a scRNA-Seq experiment

This section outlines the preparation of the four essential files required for running MINI-EX from a scRNA-Seq experiment. In the examples below, we show how to extract the required input from a scRNA-Seq experiment processed with the Seurat R package. It is important to note that all input files, except for the marker genes, should have unique row names and data should be stored in tab-delimited format without quotation marks. In the examples below, we will use the Seurat object available at https://zenodo.org/records/10423816?token=eyJhbGciOiJIUzUxMiJ9.eyJpZCI6ImUzZmQyYmRhLWUwMTItNGY1My04NDI4LTk0MThlZjRhYzE1OSIsImRhdGEiOnt9LCJyYW5kb20iOiJhZTM2ZjI2ZjA5YTQzMWQ2NjIxZDQwNTEwMzQxNTExZSJ9.Ge8ym9h2nTvFZPreJz85PBrelLaiGXh3b_CjOW9xzLod-Bi89ZVlWczrW3MlMVWGX55rOKEcP70PI8n2qegQEw.

1. **Loading the Seurat object**. In the examples below, we assume that the Seurat object has been loaded and stored in the variable ‘seurat.obj’, as demonstrated below. MINI-EX requires that all scRNA-Seq-related input files from the same dataset begin with the same prefix; we therefore store the prefix of our dataset in the variable ‘out.prefix’.

library(Seurat)
seurat.obj <-readRDS(file = "/input_dir/Wendrich.rds")
• ut.prefix <-"athWendrich"
• ut.dir <-"/output_dir"
2. **Expression matrix**: this file provides the expression levels of each gene in each cell. The format for this input file should be a tab-separated file with gene identifiers as rows and cell identifiers as columns. Specifically, the cell in row X and column Y should contain the expression level of the gene X in cell Y. While we recommend providing the original (non-normalized) expression levels (counts), alternative approaches may be considered (see Note 4). The expression matrix can be generated using the following code.

write.table(as.matrix(seurat.obj@assays$RNA@counts),
file = sprintf("%s/%s_matrix.tsv", out.dir, out.prefix),
sep = "\t", quote = FALSE)
3. **Marker genes**: this is a list of differentially expressed genes (markers) between a cluster and all other clusters in the scRNA-Seq dataset, for each cluster in the dataset. Only up-regulated markers are allowed (see Note 5) for MINI-EX. This file should follow the format of Seurat’s ‘FindAllMarkers’ function and can be generated using the code below.

markers <-FindAllMarkers(seurat.obj, only.pos = TRUE)
write.table(markers, sprintf("%s/%s_allMarkers.tsv", out.dir, out.prefix),
sep = ’\t’ , quote = FALSE)
4. **Cluster identities of cells**: this is a tab-separated file with cells in the first column and their corresponding assigned clusters in the second column.

cells2cluster <-FetchData(seurat.obj, vars = ’ident’)
write.table(cells2cluster, sprintf("%s/%s_cells2clusters.tsv", out.dir, out.prefix),
sep=’\t’, quote = FALSE, col.names = FALSE)
5. **Cluster annotation**: this is a tab-separated file with cluster identities (matching the previous file) in the first column and their annotations in the second column. The user can also add an optional third column to configure the cluster order in the regmap figure (see Section 3.8), where the column index of each cluster in the plot is added as a third column. This file can be created manually. Cluster annotations should not contain underscore characters. The first lines of the cluster annotation file for our example dataset are provided below; the full file can be downloaded from https://zenodo.org/records/10423816?token=eyJhbGciOiJIUzUxMiJ9.eyJpZCI6ImUzZmQyYmRhLWUwMTItNGY1My04NDI4LTk0MThlZjRhYzE1OSIsImRhdGEiOnt9LCJyYW5kb20iOiJhZTM2ZjI2ZjA5YTQzMWQ2NjIxZDQwNTEwMzQxNTExZSJ9.Ge8ym9h2nTvFZPreJz85PBrelLaiGXh3b_CjOW9xzLod-Bi89ZVlWczrW3MlMVWGX55rOKEcP70PI8n2qegQEw.

|  |  |
| --- | --- |
| 16 | pericycle |
| 24 | xylem |
| 9 | epidermis |
| 28 | trichoblast |
| 0 | LRC |

### 3.5. Preparing input files for non-supported species

Users planning to run MINI-EX on the supported species only need the four files related to an scRNA-Seq experiment. For non-supported species, in addition to these files, a list of TFs specific to the species must also be provided. Additionally, users may choose to include further files based on the specific MINI-EX features required for their analysis. The description of these additional files is provided below. It is important to verify that the gene identifiers used in these files match those used in the scRNA-Seq experiment.

1. **List of TFs** (mandatory): this file is essential for inferring the initial GRN as depicted in Figure 1, Step 1. This file only has one column corresponding to gene identifiers of transcription factors. The list of transcription factors of the species of interest can be downloaded from PlantTFDB (http://planttfdb.gao-lab.org/index.php) (***30***).
2. **Motif mapping** (optional): this file is necessary to execute Step 2 of MINI-EX, which involves filtering the original GRN using TF motif-supported information. Detailed instructions on generating this file can be found in (***31, 32***) (see Note 6).
3. **Transcription factor family and motifs** (optional): this is the second requirement for running Step 2 of MINI-EX. It establishes the connections between each TF, its family, and the availability of motif information. This file is tab-delimited and includes four columns:

a. TF identifier,
b. TF family,
c. ‘Y’ if motif information is available, ‘N’ otherwise,
d. Motif information

The data for the first two columns can be downloaded from PlantTFDB (http://planttfdb.gao-lab.org/index.php; (***30***), while TF motif information can be obtained from JASPAR (https://jaspar.genereg.net/downloads/) (***33***) and CisBP (http://cisbp.ccbr.utoronto.ca/bulk.php) (***34***). An example of the TF family and motifs file is provided below, with the first lines of the file available on the MINI-EX GitHub for rice. Note that for the TF on the last line, no motif information is available.

4. **GO file** (optional): this file is only required if the user has specified a list of terms of interest. It serves to prioritize regulons based on these user-defined terms during the step 4, with the last column (GO term description) used for filtering relevant GO terms. In case no terms of interest are provided, the GO file can still be provided to generate GO enrichment information about the regulons’ TGs, but this will not affect the final GRNs or regulon prioritization. The GO file lists gene-GO associations for biological processes in a tabular format, consisting of four columns:

|  |  |  |  |
| --- | --- | --- | --- |
| LOC_Os01g04750 | RAV | Y | M06692,M01648,M06693,M09748,M09749 |
| LOC_Os01g04800 | RAV | Y | M06692,M01648,M06693,M09748,M09749 |
| LOC_Os05g47650 | RAV | Y | M06692,M01648,M06693,M09748,M09749 |
| LOC_Os01g49830 | RAV | Y | M06692,M01648,M06693,M09748,M09749 |
| LOC_Os01g01430 | NAC | N |  |

a. GO term identifier,
b. gene identifier,
c. evidence code,
d. GO term description.

The contents of this file must be obtained from the data provider for the species of interest. To ensure accurate results in enrichment analysis, we recommend performing parental term propagation, which may be accomplished using various available tools (for example GOATOOLS) (***35***). For non-model species, which generally lack extensive annotations, we recommend retaining all the information. However, for more extensively studied species, it can be advantageous to filter annotations based on experimental evidence codes (the list of codes to keep can be found on the Gene Ontology website: https://geneontology.org/docs/guide-go-evidence-codes/) (see Note 3). An example is shown below using the gene-GO association file provided in the MINI-EX GitHub for rice.

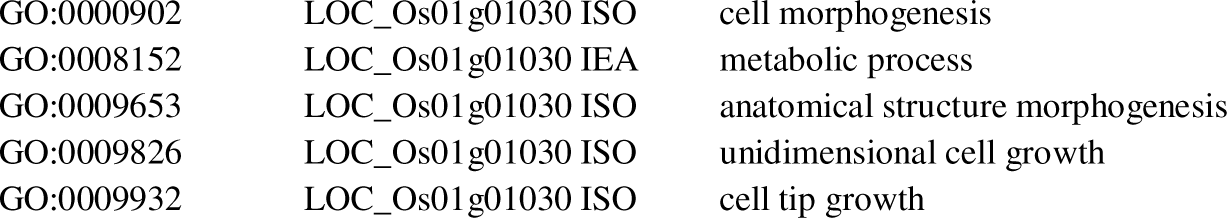
5. **List of gene aliases** (optional): this file allows users to display gene aliases instead of gene identifiers on the output files provided by MINI-EX. It is a tab-formatted file with two columns: gene identifier and gene alias. An example is shown below using the first lines of the gene alias file provided in the MINI-EX GitHub for rice. If no gene aliases are available, MINI-EX will display the original gene identifiers in the output files.

|  |  |  |
| --- | --- | --- |
| GO:0000902 | LOC_Os01g01030 ISO | cell morphogenesis |
| GO:0008152 | LOC_Os01g01030 IEA | metabolic process |
| GO:0009653 | LOC_Os01g01030 ISO | anatomical structure morphogenesis |
| GO:0009826 | LOC_Os01g01030 ISO | unidimensional cell growth |
| GO:0009932 | LOC_Os01g01030 ISO | cell tip growth |

LOC_Os07g46460 ABC1
LOC_Os07g46460 OsFd-GOGAT
LOC_Os07g46460 SPL32
LOC_Os07g46460 ES7
LOC_Os07g39750 AchE

### 3.6. Setting MINI-EX parameters

The behavior of MINI-EX depends on several parameters to be set by the user (see Note 7 on using the default parameters). Here, all parameters will be described and the rationale behind tuning those parameters provided in accompanying notes. Figure 2 illustrates how these settings are integrated in the MINI-EX algorithm.

1. **Do motif analysis**: the default value is “true”, resulting in the conventional way of running MINI-EX including a motif enrichment step (see step 2; Figure 2). Setting this parameter to “false” will omit the motif enrichment step in the algorithm. Disabling this option is not recommended because it will reduce the precision of the resulting networks (see Note 1 for more details). However, by omitting this step, no motif or TF family information is required anymore. This allows running MINI-EX for any non-supported species, without the need to generate the additional files described in Section 3.5.
2. **Terms of interest** (optional): this is an optional file that the user can provide, affecting how the regulons are ranked. This file contains a set of terms that are used to retrieve GO terms (see Note 8 on how this is done) related to the research question of the user. When this file is not provided (set to “null” in the config file), MINI-EX will use its standard procedure to rank regulons, based only on network statistics and cluster specificity (see Note 9 for more details). When the user does provide this file, MINI-EX will use GO information to give a better rank to regulons that have an overrepresentation of the GO terms of interest among their target genes (see Note 10 for more details).
3. **Motif filter**: this parameter can either have the value “TF_motifs” or “TF-F_motifs” (default) and affects how strict the motif enrichment filtering is in step 2 (see Figure 2). Setting it to “TF_motifs” results in filtering for regulons supported by the specific binding motif(s) of a regulon’s TF, whereas “TF-F_motifs” (default) results in a more liberal filtering, keeping regulons supported by any TF binding motif within the TF protein family. As a result, using “TF-F_motifs” allows keeping regulons controlled by TFs that lack motif information (see Note 11 for more details) and thus increases the number of TFs that can be studied.
4. **Top markers**: this is the number of marker genes for each cluster that are selected for the marker gene enrichment and filtering in step 3a of MINI-EX. This parameter can strongly affect the size of the final regulons with higher numbers leading to bigger regulons (see Note 12 for more details).
5. **Expression filter**: this is the percentage of cells in a single-cell cluster that shows expression of a TF, which is used as the cutoff in step 3b (see Figure 2) to filter out regulons with a TF that is not expressed in that cluster. The higher this number, the more stringent the filtering for what is considered an expressed TF (see Note 13 for more details).
6. **Top regulons**: this parameter corresponds with the top number of regulons per cluster that are displayed in the visualizations of the MINI-EX output. It solely affects what is displayed in the output figures. The GRNs and ranking remain the same.

### 3.7. Running the pipeline

Running MINI-EX involves setting up the config file (‘miniex.config’) with all the necessary input paths and parameters and executing the following command:

nextflow -C miniex.config run miniex.nf

MINI-EX enables the concurrent processing of multiple scRNA-Seq datasets. This can be achieved by incorporating the wildcard character ‘*’ in the names of the four mandatory input files, as explained in Section 3.3. For example, using ‘*_allMarkers.tsv’ will find all files ending with ‘_allMarkers.tsv’ but starting with different dataset names (i.e. different prefixes). Please note that this feature only allows running MINI-EX on multiple datasets of the same species, as the remaining files and parameters, including the terms of interest, remain fixed in the config file.

MINI-EX is primarily designed for HPC clusters and is set to run with the job scheduler SLURM by default:

executor {
name = ’slurm’ queueSize = 5
}

Before initiating MINI-EX, users should verify which job scheduler (executor) is available on their system and adjust the ‘name’ parameter within the ‘executor’ scope accordingly (see Note 14). If the user does not intend to run MINI-EX on an HPC cluster, they can set the ‘name’ parameter to ‘local’. The list of acceptable values for this parameter can be found in the Nextflow documentation: https://www.nextflow.io/docs/latest/executor.html.

As shown above, the default value for the ‘queueSize’ parameter is five, meaning that Nextflow is only allowed to run five tasks in parallel. This value can be adjusted by the user as needed, and must be set to zero if the user does not want to set a limit.

Depending on the user’s dataset, some steps may require more or less resources. Default memory requirements are specified in the config file for each process; however, for some datasets they may not be sufficient. If MINI-EX fails at a particular step either with a “MemoryError” or without any specific error message provided by MINI-EX, it is highly probable that the process was killed due to insufficient memory. This can be corrected by increasing the ‘memory’ parameter of the corresponding dataset in the config file:

withName: run_grnboost {
memory = ’20 GB’
cpus = 5
}

As constructing the GRN (Step 1), performed by GRNBoost2 (***26***), is one of the most resource-intensive steps, it is possible to store the GRNBoost2 output for reuse in subsequent runs on the same dataset. This can be accomplished using the ‘grnboostOut’ parameter. For the first run, this parameter must be set to ‘null’, indicating that the base GRN needs to be computed. For subsequent runs, this parameter can be set to the path of the GRNBoost output file:

grnboostOut = "/$baseDir/example/OUTPUTS/GRNBoost2_output/*_grnboost2.tsv"

### 3.8. The MINI-EX output files

When MINI-EX has been run successfully, output files are stored in separate folders, named GRNBoost2_output, GOenrichment_output, regulons_output and figures. In this section, the different output files contained in these folders will be described along with their interpretation. We first discuss the ranked regulons file and the regulons file, as these can be considered the two most important output files of MINI-EX, followed by a discussion of the output figures.

1. **Ranked regulons file** (*_rankedRegulons.xlsx or *_rankedRegulons.tsv): this file, provided in different file formats for user convenience, contains metadata and statistics on the final regulons inferred by MINI-EX, including their relevance ranking.

a. **TF information** (columns 1-5; Table 1): column 1 contains the gene ID of the TF, column 2 its alias (i.e. gene symbol), column 3 shows whether the TF is already known (GO information is available for it) and whether it is a relevant TF (at least one of the associated GO terms matches a term of interest given by the user; see Section 3.5 and Note 8. Columns 4 and 5 show the GO terms associated with the TF and their description, respectively.
b. **Cluster-related information** (columns 6-9; Table 2): column 6 contains the cluster label and column 7 the annotation of that cluster (as given in the cluster annotation input file). Column 8 shows whether the TF of the regulon is a marker gene for that cluster, and column 9 gives the total number of regulons that were retained for that cluster.
c. **Regulon importance metrics** (columns 10-14; Table 3): column 10 contains the number of TGs in the regulon. Column 11 contains the cluster specificity, which corresponds to the q-value of enrichment between the regulons’ TGs and the marker genes of that cluster (calculated in step 3a of MINI-EX; see Figure 2). Columns 12-14 contain three network centrality metrics: out-degree, closeness and betweenness, which measure how central the TF is embedded within the network structure (see Note 15).
d. **Enrichment of TGs for GO terms of interest** (columns 15-18; Table 4): these four columns are only present when the input file with terms of interest was provided by the user. The column “GO_enrich_qval” contains the q-value of the most significantly enriched GO term of interest among the TGs in that regulon, “GO_enrich_term” contains the corresponding GO term, “GO_enrich_desc” the description of that GO term and the column “#TGs_withGO” the number of TGs in the regulon that are associated with that GO term.
e. **Regulon ranking** (Table 5): the column “borda_rank” shows the global Borda rank over all cluster-specific networks, while “borda_clusterRank” gives the Borda rank of the regulon within the network of one single-cell cluster (see Note 16). The Borda rank utilizes the cluster specificity, network centrality of the TF, and optionally also the enrichment of a regulon’s TGs for GO terms of interest to produce a final ranking of the regulons based on the most relevant regulon importance metrics (see Note 10).
2. **Regulons file** (*_regulons.tsv): this file contains the final MINI-EX network, with each line representing a regulon. It is a tab-delimited file with the TF of that regulon as the first column, the regulon’s single-cell cluster as the second column, and a comma-separated list of the regulon’s target genes as the last column.

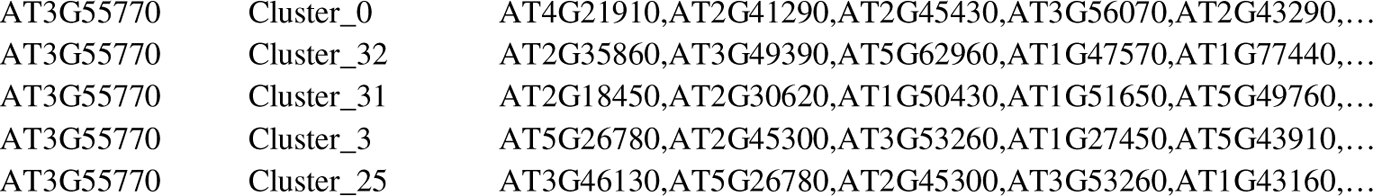

**Table 1.** Ranked regulons result table, columns related to TF information.

| TF | alias | hasTFrelevantGOTerm | GOTerm | GODescription |
| --- | --- | --- | --- | --- |
| AT3G58710 | WRKY69/<br>ATWRKY69/<br>ABT4 | unknown_TF | - | - |
| AT5G26660 | MYB86/<br>ATMYB4-old/<br>ATMYB86/<br>ATMYB4 | known_TF | GO:0006139,G<br>O:0006351,... | nucleobase-containing compound<br>metabolic process,DNA-templated<br>transcription,... |
| AT5G16600 | MYB43/<br>AtMYB43 | known_TF | GO:0006082,G<br>O:0006139,... | organic acid metabolic<br>process,nucleobase-containing<br>compound metabolic process,... |
| AT3G61850 | DAG1 | known_TF | GO:0006139,G<br>O:0006351,... | nucleobase-containing compound<br>metabolic process,DNA-templated<br>transcription,... |
| AT5G18270 | ANAC087 | unknown_TF | - | - |

**Table 2.**
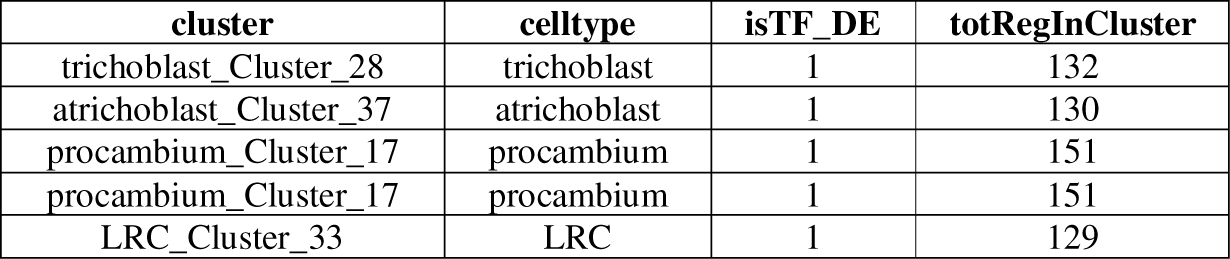
Ranked regulons result table, columns with cluster-related information.

**Table 3.** Ranked regulons result table, columns with regulon statistics for ranking.

| #TGs | qval_cluster | out-degree | closeness | betweenness |
| --- | --- | --- | --- | --- |
| 673 | 1,4E-254 | 0,821734 | 0,126227 | 0,006404 |
| 668 | 0 | 0,817625 | 0,131409 | 0,005853 |
| 683 | 8,8E-267 | 0,825877 | 0,147863 | 0,00381 |
| 683 | 7,6E-157 | 0,825877 | 0,147863 | 0,003663 |
| 694 | 0 | 0,864259 | 0,136936 | 0,002872 |

**Table 4.**
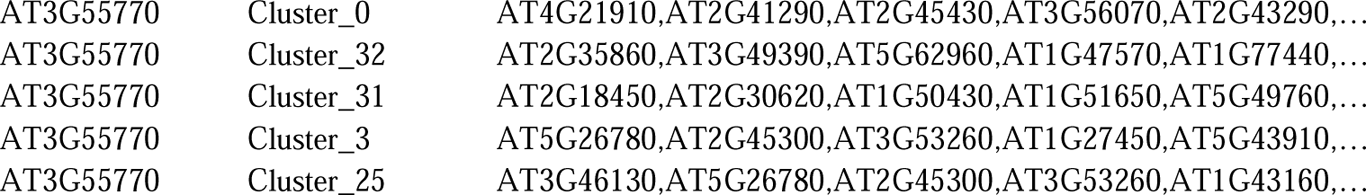
Ranked regulons result table, columns with regulon gene ontology enrichment information.

**Table 5.** Ranked regulons result table, columns with regulon ranking information.
3. **Edge table file** (“*_edgeTable.tsv”): this file (Table 6) combines the most important information about the network as a tab-separated edge list. Each line represents and edge in the network, with in the first column the gene ID of the TF, the second column the gene ID of the TG, the third the cluster ID, the fourth the global Borda rank of the regulon, the fifth the cluster-specific Borda rank, and the sixth is the GRNBoost2 evidence score of that edge, which is a measure for the degree of coexpression.
4. **TF info file** (“*_TF_info.tsv”): this file (Table 7) shows detailed information about how TFs in the initial TF list are filtered during the different steps in MINI-EX. The TF info file is a table with a TF on each row. The first column shows whether the TF is found as expressed in the count matrix, the second shows whether it is part of a regulon in the GRNBoost2 GRN, the third whether it is retained as a regulon after motif mapping filtering in step 2 (see Figure 2), and the fourth shows if the TF is part of the final networks after expression filtering in step 3. In case the TF is expressed (included in the expression matrix), all following columns show the percentage of cells in which the TF is expressed for each single-cell cluster

| borda_rank | borda_clusterRank |
| --- | --- |
| 1 | 1 |
| 2 | 1 |
| 3 | 1 |
| 4 | 2 |
| 5 | 2 |

**Table 6.** Edge table result file.

| TF | TG | cluster | borda_rank | borda_clusterRank | weight |
| --- | --- | --- | --- | --- | --- |
| AT1G01010 | AT2G32240 | Cluster_11 | 4584 | 183 | 1,39859E+16 |
| AT1G01010 | AT4G36990 | Cluster_11 | 4584 | 183 | 7,85487E+14 |
| AT1G01010 | AT1G23440 | Cluster_11 | 4584 | 183 | 4,33944E+15 |
| AT1G01010 | AT1G73260 | Cluster_11 | 4584 | 183 | 4,18308E+15 |
| AT1G01010 | AT4G01070 | Cluster_11 | 4584 | 183 | 3,77707E+15 |

**Table 7.** Transcription factor information table.
5. **GRNBoost2 network** (“*_grnboost2.tsv”): this file contains the GRN inferred by GRNBoost2. Each line in this file corresponds to a directed edge in this network, with the first column being the gene ID of a TF and the second column the gene ID of its TGs. The third column is the evidence score calculated by GRNBoost2 for this edge.

|  | isTF_expressed | isTF_expressionR<br>egulons | isTF_TFBS<br>Regulons | isTF_final<br>Regulons | LRC-0 | LRC-1 | ... |
| --- | --- | --- | --- | --- | --- | --- | --- |
| AT1G01010 | 1 | 1 | 1 | 1 | 1.4 | 1.5 | ... |
| AT1G01030 | 1 | 1 | 0 | 0 | 2.3 | 0.5 | ... |
| AT1G01060 | 1 | 1 | 1 | 1 | 0.8 | 1.2 | ... |
| AT1G01250 | 0 | 0 | 0 | 0 |  |  | ... |
| AT1G01260 | 1 | 1 | 1 | 1 | 10.1 | 15.5 | ... |

6. **GO enrichment of regulons** (“*_enricherGO.txt”): this file (Table 8 and 9) contains the raw output of GO enrichment on the TGs of each regulon. The first column contains the gene set ID, denoting which set of genes (here the TGs of a regulon) was tested for enrichment. The set ID is constructed as the regulon’s TF gene ID, followed by the cluster name of that regulon, joined by an underscore (e.g. AT3G44750_Cluster_4). The second column is the enriched GO term and the third is the p-value of the enrichment. The fourth column contains the q-value (adjusted p-value by using the Benjamini-Hochberg procedure (***36***)) and the fifth column the enrichment fold (the ratio between the observed fraction of hits in the gene set and the fraction in the background). The sixth column shows the size of the gene set (i.e. number of TGs in the regulon), the seventh column shows the feature set size (i.e. the total number of genes associated with the GO term) and the eighth column shows the information content (IC) (***37***) of the GO term, which is a metric that ranges between 0 and 1 that describes how general (low values) or specific (high values) the GO term is. The ninth column shows the number of genes that the gene set and feature set have in common, and the last column displays those genes as a comma-separated list of gene IDs.
7. **Regmap figure** (“*_regmap_*.svg”): this figure (Figure 3) is very informative to identify the most relevant TFs in different clusters. Several regmap figures are generated, each visualizing an increasing number (as indicated in the file name; e.g. *_regmap_25.svg) of the (e.g. 25) top most important regulators per cluster. The figure shows a grid in which the rows correspond with TFs and the columns with different single-cell clusters. Columns are also annotated by their cell type identity, as given in the cell cluster annotation input file, and if an optional index column is added to this file (see Section 3.4), the specified column index is used in the plot. In each slot, a circle is plotted where the size of the circle corresponds to the maximum expression level (average of three highest expressing cells) of the TF in the given cluster and the shade of the color corresponds to the cluster-specific Borda rank. This plot allows one to visually identify the most important regulators in a given cell type, according to the Borda ranking procedure. One can also easily identify shared regulators, active in multiple cell types. If the clusters follow a known developmental trajectory, patterns of increasing or diminishing regulator importance can be tracked over that trajectory.
8. **Clustermap figure** (“*_clustermap.svg”): this figure (Figure 4) shows a clustered heatmap of the regulon size in different clusters. The x-axis represents the different single-cell clusters and the y-axis the different TFs that can regulate different sets of target genes in different clusters. The heatmap shows the number of TGs in the regulon of the given cluster. This plot visualizes similarities between single-cell clusters based on their regulons.
9. **DE and specificity heatmap figures** (“*_heatmapDEcalls.svg” and “*_heatmapSpecificity.svg”). Both figures show the different single-cell clusters on the x-axis and the different TFs on the y-axis. The TFs are annotated with green if they are a relevant TF (i.e. they match a GO term of interest), with yellow if they are a known TF (annotated to any non-relevant GO term) or with gray if they are not annotated to a GO term. For the DE heatmap (Figure 5), the colors in the heatmap indicate whether the TF is upregulated (blue), expressed in at least 10% of the cells of that cluster (white) or expressed in less than 10% of the cells of that cluster (gray). For the specificity heatmap figure the negative logarithm of the cluster specificity q-values (see cluster specificity column in ranked regulons file) is shown. Both files visualize where TFs are expressed in a cluster-specific manner.
10. **Log file** (*_log.txt). This file contains a summary of the input files and parameters used in a given MINI-EX run, as well as summary statistics on the initial data and the inferred GRNs. Number of regulons and median regulon size are given after each filtering step in MINI-EX, allowing to identify the impact of each filtering step. Finally, the log file shows the regulon importance metrics and ranking procedure (see Note 9 and 10) used for the Borda ranking of regulons in step 4 (see Figure 2).

|  |  |  |
| --- | --- | --- |
| AT3G44750 | AT4G12600 | 228.43688171026278 |
| AT1G63100 | AT1G50490 | 226.98249498720028 |
| AT3G44750 | AT5G27120 | 207.9133763122063 |
| AT1G63100 | AT3G51720 | 206.86956547679816 |
| AT3G44750 | AT4G25630 | 204.25228196977886 |

**Figure 3.**
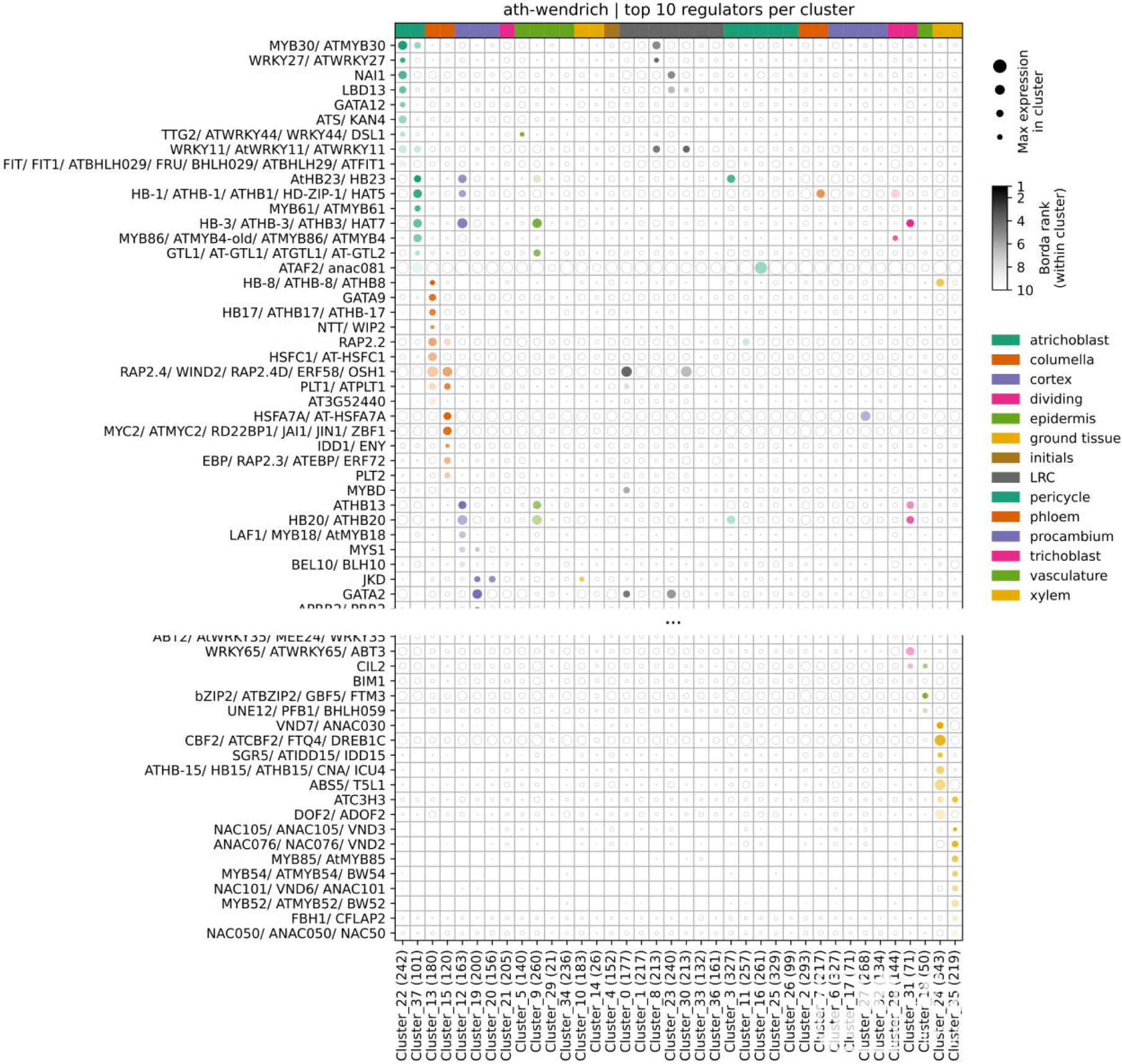
Regmap figure of the Wendrich *et al.* (2020) results. The top 10 most important regulators of each cluster (based on the cluster-specific Borda rank) are selected. Gene aliases are shown on the y-axis and clusters on the x-axis. For each cluster-specific network, the total number of regulons is added after the cluster name. The shade of the color of each circle corresponds to the cluster-specific Borda ranking of the given regulon while the size corresponds to the maximum expression level of the regulator within that cluster.

**Figure 4.**
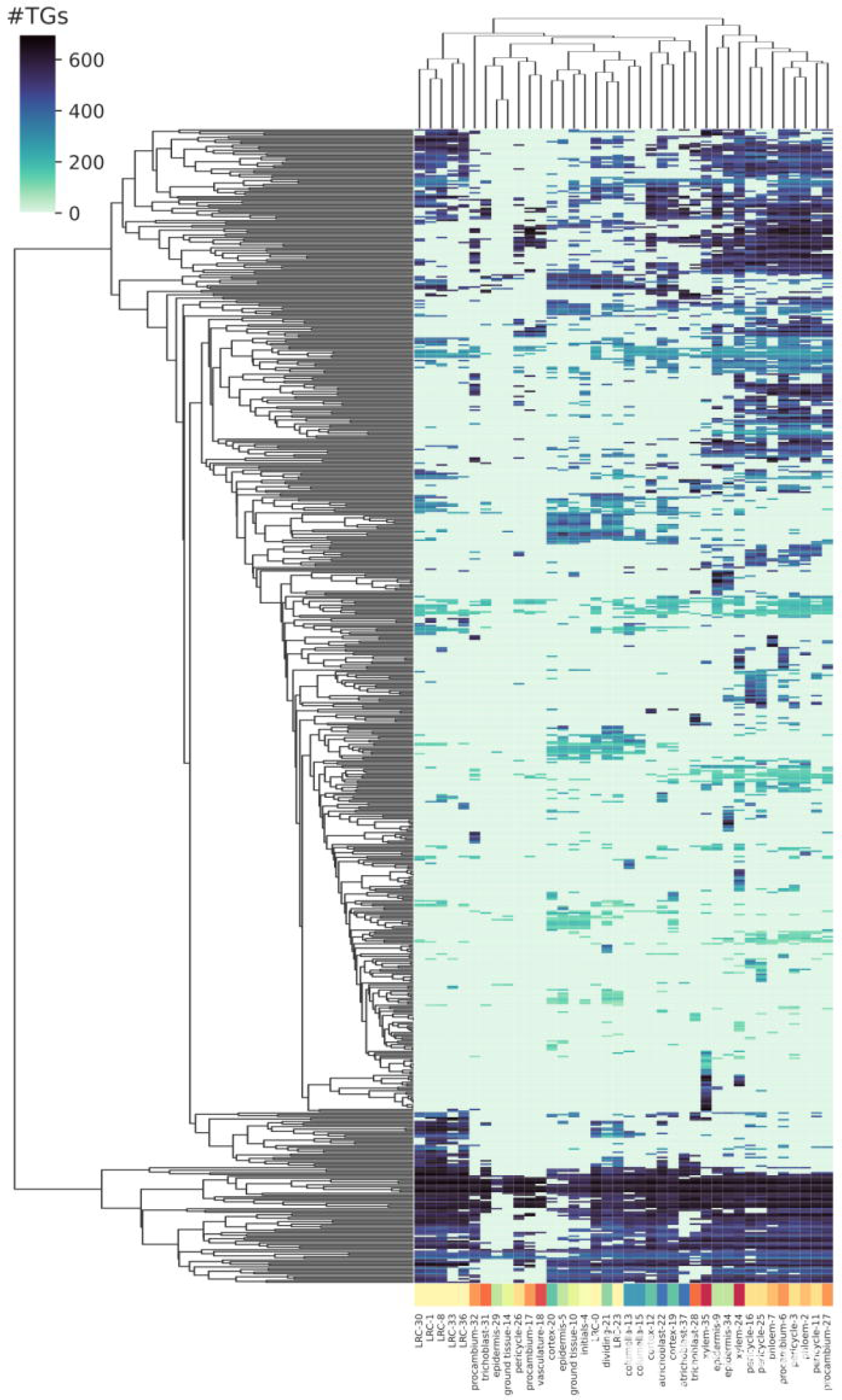
Clustermap figure of the Wendrich *et al.* (2020) results. This figure shows a clustered heatmap of the regulon size. The x-axis represents single-cell clusters and the y-axis individual regulons. The color of the heatmap represents the number of target genes the regulon has.

**Figure 5.**
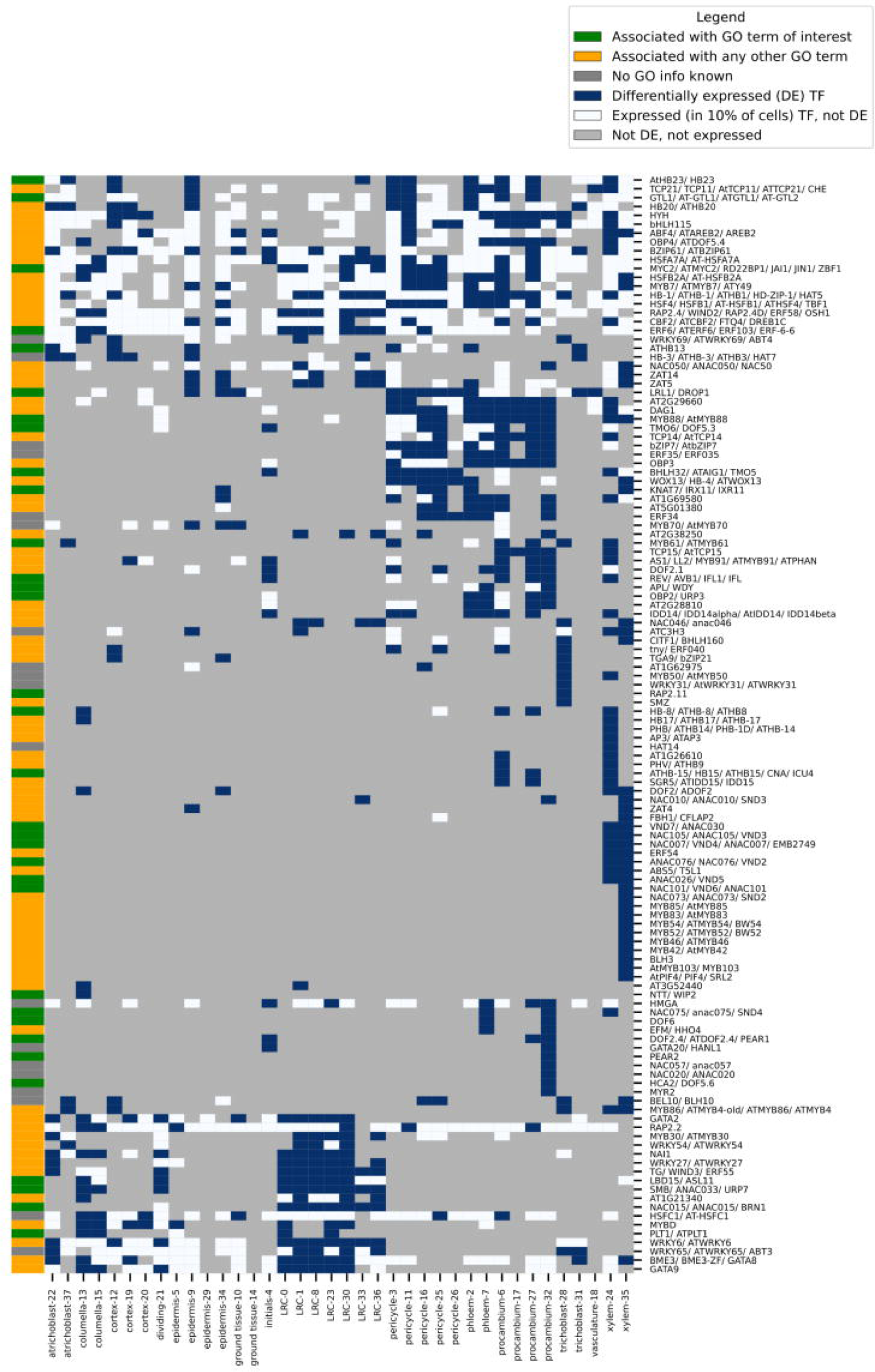
DE heatmap figure of the Wendrich *et al.* (2020) results. This figure shows a heatmap with the differentially expressed (DE) transcription factors (TFs) in at least one single-cell cluster. The x-axis represents single-cell clusters and the y-axis individual TFs. The color of the heatmap is blue if the TF is DE (= marker gene) in the cluster, white if it is not DE but it is expressed in more than 10% of cells, and gray if it is neither DE nor expressed in more than 10% of cells in the cluster. TFs are annotated as green if they are associated with a GO term of interest, yellow if they are associated with any other GO term and gray if they are not associated with a GO term.

**Table 8.** Gene ontology enrichment output file.

| #set_id | ftr_id | p-val | q-val | enr_fold |
| --- | --- | --- | --- | --- |
| AT1G14900_Cluster_32 | GO:0000003 | 0.000563632 | 0.00224408 | 2.56505 |
| AT1G14900_Cluster_32 | GO:0003006 | 0.00186918 | 0.00596871 | 2.45494 |
| AT1G14900_Cluster_32 | GO:0006139 | 0.00268205 | 0.0079436 | 2.08545 |
| AT1G14900_Cluster_32 | GO:0006351 | 0.000607938 | 0.00238854 | 2.64713 |
| AT1G14900_Cluster_32 | GO:0006355 | 0.000541584 | 0.00217134 | 2.67711 |

**Table 9.** Gene ontology enrichment output file.

| set_size | ftr_size | ftr_ic | n_hits | hits |
| --- | --- | --- | --- | --- |
| 202 | 688 | 0.28924 | 16 | AT5G61230,AT1G72150,AT4G30520,... |
| 202 | 629 | 0.298994 | 14 | AT1G72150,AT4G30520,AT2G32280,... |
| 202 | 952 | 0.253911 | 18 | AT3G61850,AT4G29230,AT5G60850,... |
| 202 | 625 | 0.299688 | 15 | AT3G61850,AT4G29230,AT5G60850,... |
| 202 | 618 | 0.300913 | 15 | AT3G61850,AT4G29230,AT5G60850,... |

### 3.9. Interpreting the results

Finally, to illustrate how the MINI-EX results can be used to extract interesting biological knowledge, we will here look more in depth at the results of two of the most informative output files: the ranked regulons file and the regmap figure. As an example, we focus on the well-studied xylem tissue, to see what information regarding its regulation is captured by MINI-EX.

First, we can visually inspect the regmap figure to identify the most important regulators. The xylem tissue consists of clusters 24 and 35 which correspond to the last two columns in the figure. At the lower part of the regmap figure, many known xylem differentiation regulators can be observed such as MYB52, MYB54, MYB85, VND2, VND3, VND6 and VND7 (***38***). The regmap shown in Figure 3 is the smallest of several generated regmap figures, only coloring the top 10 best ranked regulators per cluster. While this regmap is the most readable, potential relevant regulators can be missed in this regmap, for example the known xylem regulators MYB46 and MYB83, respectively ranked 18^th^ and 20^th^ in xylem cluster 35. Similarly, correspondence between clusters can be missed in the smaller regmaps. This is the case for VND7 in Figure 3, which is the top regulator for xylem cluster 24 but is not shown to be relevant for xylem cluster 35. However, looking at the ranked regulons file, we can see that VND7 is still ranked 13^th^, indicating a relevant role for this regulator in both clusters.

Another putative xylem regulator HB-8 can be seen higher on the plot. HB-8 is also a known regulator involved in xylem formation (***39***). It is shown higher up in the figure because it is also a well-ranked regulator in the columella tissue, and columella regulators are (arbitrarily) plotted prior to xylem regulators. While HB-8 has indeed been confirmed to show increased expression not only in the xylem but also in the columella, a potential role for this gene as a regulator in columella differentiation has not been described, indicating a potential new discovery by MINI-EX. While both tissues are derived from the same stem-cell niche, they are not considered very similar tissues. This shared regulator suggests shared downstream effects in both xylem and columella. However, if we extract the TGs of HB-8 in the xylem and columella from the regulons file (*_regulons.tsv), we see that out of 672 TGs in xylem and 526 TGs in columella, only 27 TGs are shared. This suggests that, besides BH-8 possibly being a regulator in both tissues, the BH-8 downstream signaling is rewired. Based on these results, it would be interesting to experimentally investigate the role of HB-8 in the differentiation of the columella, for example by performing a columella-specific knock-out experiment.

In conclusion, this example shows that MINI-EX is a valuable tool to predict known and novel regulators in different tissues and processes of interest, as well as to discover new functions of known regulators. The interpretation of MINI-EX’s summarizing output files and figures can offer researchers profound insights, enabling them to formulate new hypotheses about the biological function of regulators and their downstream effects.

## 4. Notes

1. In step 2, filtering is performed on the regulons of the initial regulatory network in two ways. First, a regulon is only kept if the TF motif is enriched in its TGs (i.e., more TGs than expected by chance have the TF binding motif). Second, for the remaining regulons, only TGs that have the TF motif are retained. As a result, regulons under the control of a TF lacking information on their binding motif are removed. While MINI-EX by default extends motifs to all family members of that TF protein family (see “motif filter” parameter; Section 3.6), increasing the number of TFs with motif information, still a fraction of TFs (around 20% in *A. thaliana*) lacks this information which causes them to be filtered out during this step. Thus, even when extending motifs to the TF family, it is important to take into account that when little motif information is available, this can result in a filtering in step 2 that is too severe. If one notices that too many relevant regulons are removed during step 2 due to lack of motif information, it might be beneficial to omit step 2 (see “do motif analysis” parameter; Section 3.6). However, for species (e.g., supported species) with sufficient TF motif information, it is recommended to include the motif mapping step as it has been shown to improve precision of the resulting GRNs (***19, 40, 41***).
2. In MINI-EX version 2.0, the way in which enrichment statistics were calculated was modified for all enrichment steps within MINI-EX (see steps 2, 3a and 4a in Figure 2). A first modification is to perform enrichment using a background model of only expressed genes (***27, 28***). This is implemented by first inferring the set of expressed genes based on the gene x cell count matrix given as input. Then, the motif mapping file, the marker gene file, and the GO file, used in the three enrichment steps of MINI-EX, are filtered to only keep expressed genes. Lastly, the filtered files are used and the set of all expressed genes is used as a background model when calculating enrichment statistics. A second modification was also made so that no enrichment statistics were calculated if there is only one hit in the gene set.
3. MINI-EX provides gene-GO association files for supported species, derived from PLAZA (***29***) and combined with GO information from TAIR (www.arabidopsis.org) (***42***) for *A. thaliana*. Originally, in MINI-EX, these contained only GO terms with experimental evidence codes (EXP, IMP, IDA, IPI, IGI, IEP, TAS, NAS, IC). Since version 2.1, all evidence codes including computational evidence codes are included for all supported species except *A. thaliana*, in order to cover a sufficient number of genes to calculate GO enrichment statistics. Given that for the model species *A. thaliana*, sufficient experimental GO information is available, a more stringent selection for high-quality (i.e. experimental) terms is opted for (since version 2.2 this also includes the high throughput experiment code HTP, but not the non-traceable author statement code NAS).
4. The expression matrix provided to GRNBoost2 as input should be a count matrix. Due to the non-linear tree-based approach GRNBoost2 takes, normalization of this matrix is possible, but not necessary (***43***). It is important to note that normalization procedures such as log-transformation or transcripts per million can introduce artifacts and should therefore not be applied (***44***). One possible normalization method is SCT normalization, as implemented by the R package Seurat. With this method, Pearson residuals are reverse-transformed into a count matrix that is adjusted for sequencing depth and corresponds to the “counts” matrix of the SCT slot in a Seurat object. Such an adjusted count matrix can be provided to MINI-EX as input. While such normalization procedures can help to reduce some biases, they are not necessary. Therefore, we recommend using unnormalized, raw counts to new users as a valid first option.
5. In step 3a (see Figure 2), MINI-EX uses a set of differentially expressed genes (DEGs; i.e., marker genes) per cluster. These DEGs should be upregulated genes (set by specifying the “only.pos = TRUE” parameter in the Seurat function FindAllMarkers). MINI-EX, as it is currently implemented (version 2.2) is only intended to infer upregulatory relationships between TF and TG. Hence, only upregulated DEGs should be given as input.
6. The motif mapping file was generated by mapping known TF binding site (TFBS) motifs to the regulatory region around all genes in the genome. If the TFBS motif of a TF is found around the gene, the motif-TG pair is added to the motif mapping file. As the aim of this chapter is to be a practical guide on how to use MINI-EX, we will not give an extensive description on the procedures used to generate the precomputed motif mapping files of which MINI-EX makes use. More detail regarding the way this file was created can be found in (***19, 31, 32***).
7. The default parameters of MINI-EX are based on benchmarking results on *A. thaliana* described in the original MINI-EX publication. We recommend evaluating how well these parameters perform, especially when using MINI-EX for species other than *A. thaliana*, using a relevant gold standard reporting TF - TG interactions or a set of known TFs for the process of interest (***19***).
8. The “Terms of interest” file contains words that can be linked to specific GO terms of interest. Using these words, MINI-EX retrieves the GO terms that contain at least one of the provided words in their description. For example, adding the term “trichome” in the Terms of interest file will result in several GO terms being selected such as “trichome branching”, “regulation of trichome morphogenesis”, and others. We encourage users to provide synonyms and related terms such as “xylem”, “vessel” and “tracheary” in order to cover all relevant GO terms. Also, when one has a broad research question such as interest in all root related processes, we suggest providing not only “root” as a term but also more specific tissues or processes such as “trichoblast”, “columella”, etc.
9. When no Terms of interest file is provided or no GO information is available, MINI-EX uses its default ranking procedure, also referred to as the standard or “std” procedure. This procedure ranks all inferred regulons based on four regulon importance metrics. These are the out-degree, closeness, betweenness (three network centrality metrics) and cluster specificity. Cluster specificity corresponds with the q-value of enrichment between cluster-specific marker genes and the TGs of a regulon (i.e. how specific are the TGs of the regulon for the cluster) calculated during step 3a (see Figure 2). These four regulon importance metrics are integrated using unweighted Borda counts. Here, the regulons are ranked on each of the metrics and the geometric mean of those four ranks is taken for each regulon. The resulting number is used to make the final ranking.
10. When both GO information is provided and the user specified a set of terms of interest, MINI-EX will use this information as reference to tailor its ranking procedure to the research question of the user, defined by these specific terms. This ranking procedure is referred to as the reference of “ref” procedure. It uses the same regulon importance metrics as the standard procedure (out-degree, closeness, betweenness and cluster specificity), with the addition of GO enrichment q-value between the TGs of a regulon and GO terms of interest (see also Note 3 and 8). The reference procedure makes use of weighted Borda counts to integrate regulon importance metrics and automatically selects which metrics to include for this. Giving weights and selecting regulon importance metrics is done based on R50 values. The R50 value is the rank at which 50% of all relevant regulons (i.e., regulons with a TF that is associated with a GO term of interest) are retrieved. Based on the ranking on each individual metric, an R50 value is calculated for each metric. Then, ranks for each metric are divided by the R50 of that metric and subsequently summed to get the weighted Borda score and final ranking. As such, metrics that perform worse at ranking relevant TFs and thus have a high R50, will contribute less to the final ranking. Finally, not all five regulon importance metrics are necessarily used to calculate the final Borda rank. MINI-EX will calculate a weighted Borda rank as described above for each possible combination of at least one regulon importance metric and select the optimal combination of metrics as the one that results in weighted Borda ranks with the best R50 value, which are used as the final ranking in the MINI-EX output. The selected combination of metrics can be found in the MINI-EX log file (since version 2.2).
11. The use of the TF family motif extension strategy aims to increase the recall of the results and is based on the observation that the binding motifs of different members of a TF protein family are often very similar due to their shared ancestry. Thus, to prevent filtering out many regulons only due to their TFs lacking motif information, this approach uses the known binding motifs of all family members as a proxy, thereby covering more TFs. However, while this strategy has been shown to increase recall, it may reduce precision because when TFs with motif information available are expanded to the family level, more spurious matches can be found that introduce noise, giving rise to more false positive TF-TG interactions in the final GRNs.
12. In step 3a, cluster-specific marker genes (i.e., differentially expressed genes calculated for each single-cell cluster) are used for enrichment and filtering (see Figure 2). In this step, regulons with TGs that are not enriched for the top N (the “top markers” parameter) most significant marker genes for each cluster are filtered out. For the remaining regulons, only TGs that are in the cluster-specific marker genes set are retained. Therefore, changing the “top markers” parameter strongly affects overall regulon sizes.
13. The “expression filter” parameter corresponds to the percentage of cells in a cluster that has to express a TF, in order for that TF to be considered expressed in that cluster. This metric does not take into account the expression level of a gene, which means that if the TF has one UMI count in a cell, the TF is considered expressed in that cell. As a result, the use of count normalization methods that can impute counts when the raw count was zero, can result in a too liberal filtering in step 3b. Conversely, on datasets with relatively low coverage, the high number of zero counts can result in too stringent filtering when using the default. Thus, as the optimal value for this parameter depends on your dataset and preprocessing steps, it is recommended to assess how many regulons are filtered out during this step (by looking at the log file and “TF info file”) and adjust this parameter accordingly. An example of such adjustment can be found in Ferrari *et al*. (2022) when running MINI-EX on maize.
14. If the user attempts to execute MINI-EX on an HPC cluster with an executor other than SLURM without modifying the ‘name’ parameter within the ‘executor’ scope, MINI-EX will result in failure with the following error message: ‘cannot run program “sbatch”’ (where ‘sbatch’ is the command used by SLURM to submit jobs).
15. In MINI-EX, three network centrality metrics are calculated (out-degree, betweenness and closeness), as implemented by the python library NetworkX (version 2.5.1) (***45***). The out-degree centrality corresponds to the number of outgoing edges of a TF (i.e., the number of TGs), divided by the largest possible out-degree, which is the number of other nodes in the network of that cluster. The betweenness of a TF is the fraction of shortest paths between any two nodes (excluding the TF) containing that TF. Closeness is the number of all other reachable nodes from a TF, divided by the average path length of a TF to all other nodes in the network. These metrics are complementary ways to estimate the importance of a TF in the network structure. Note that each of these metrics is calculated for the cluster-specific GRN the regulon is part of, independent of the other cluster-specific GRNs.
16. The Borda rank is calculated both in a global way and in a cluster-specific manner. For the global ranking, the regulons of all cluster-specific networks are taken into account when obtaining the ranks based on the different regulon importance metrics (which are themselves calculated within the cluster-specific GRN of that regulon; see Note 15), often including regulons under the control of the same TF, but in a different single-cell cluster with different TGs. This is opposed to the cluster-specific ranking that makes use of the same regulon importance metric values as the global ranking, but only creates a ranking of regulons within one cluster. Based on the ranks on each individual metric, the Borda rank is calculated in the same way for both the global and cluster-specific ranking.

## Acknowledgments

This work was supported by an Industrieel Onderzoeksfonds grant from Ghent University (F2020/IOF-StarTT/151; IOF.PRO.2021.0017) to S. L. and K. V., and a Bijzonder Ondersoeksfonds grant from Ghent University (BOF24Y2019001901) to N. M. and The Research Foundation - Flanders (FWO; Odysseus II G0D0515N) to J.S. We thank Herman De Beukelaer for helping us to improve enrichment analysis and Camilla Ferrari for developing the initial version of MINI-EX.

## Notes

https://github.com/VIB-PSB/MINI-EX

### Competing Interest Statement

The authors have declared no competing interest.

